# Digoxin Inhibits BaxΔ2-induced Neuronal Cell Death

**DOI:** 10.64898/2026.03.16.712124

**Authors:** Qi Yao, Jennifer M. Sorescu, Ishita Nahush Amin, Alexander Julian, Jennifer Heo, Dominique Philoctete, David Minh, Jialing Xiang

## Abstract

Pro-death Bax isoform BaxΔ2 forms protein aggregates in Alzheimer’s neurons, triggering stress granule formation and neuronal cell death. In seeking chemical ligands to prevent BaxΔ2 monomer aggregation, we performed *in silico* screening of FDA-approved drugs using computational docking. This screening identified a group of compounds that bind to the hydrophobic pocket of BaxΔ2. Subsequent wet-lab testing revealed that digoxin could block neuronal cell death at nanomolar concentrations (50 to 100 nM). Importantly, digoxin’s protective role is specific to BaxΔ2-induced cell death and is independent of its primary cardio-action on Na/K-ATPase. Further investigation suggests that digoxin does not significantly affect the formation of BaxΔ2 aggregates but may instead modulate BaxΔ2 protein levels. Although the therapeutic use of digoxin for Alzheimer’s disease is not feasible due to its narrow therapeutic window and toxicity, these findings open the door for chemical modification of digoxin, or development of similar compounds, to prevent BaxΔ2-mediated neuronal cell death in Alzheimer’s disease.

## 1. Introduction

BaxΔ2 is an alternative splicing isoform of the pro-death Bax, which has been extensively studied in both anti-cancer therapy and autoimmune diseases (1–3). However, the apoptotic function of the Bax family in the brain remains inconsistent and debatable. We recently found high levels of BaxΔ2 protein accumulated in Alzheimer’s disease (AD)-susceptible regions, such as the hippocampus and frontal lobe (4, 5). Up to 30% of AD neurons had detectable BaxΔ2 protein aggregates. Further studies in neuronal cells revealed that BaxΔ2 aggregation triggers the formation of stress granules (SGs), and the interaction between SGs and the C-terminal of BaxΔ2 leads to cell death (4). There is, however, no evidence of physical interaction between BaxΔ2 aggregates and Tau tangles (4).

Compared to Baxα, BaxΔ2 lacks one of six exons due to alternative splicing. In Baxα, helix α1 -- primarily encoded by exon 2 -- functions as a mitochondrial targeting signal and a cover for the hydrophobic pocket formed by helices α2-5, encoded mostly by exons 3-5 (6–8). The helix α9, encoded by exon 6, acts as a transmembrane domain (9). Upon receiving a death stimulus, Baxα undergoes a confirmational change that exposes the hydrophobic pocket, enabling homo-oligomerization and the formation of ring pores on the mitochondrial outer membrane (9). Unlike Baxα, BaxΔ2 lacks helix α1, impairing its ability to target mitochondria and permanently exposes its hydrophobic pocket, resulting in aggregation in the cytosol without a death stimulus (3). Subsequently, these oligomerized BaxΔ2 proteins are accumulated in the cytosol as large protein aggregates and trigger cell death (10).

In this study, we conducted computational screening of chemical ligands from FDA-approved drugs against the BaxΔ2 monomer. After identifying a group of candidates *in silico*, we performed *in cell* assays using a neuronal cell culture system to assess their cytotoxicity, and ability to prevent BaxΔ2-induced cell death and BaxΔ2 protein aggregation.

## 2. Materials and methods

### 2.1. Virtual Drug Screening

Potential drugs preventing BaxΔ2 homo-oligomerization were screened using Autodock Vina (11). A previously simulated BaxΔ2 structure (12) was used as the receptor, and the “Drugbank FDA only” ZINC15 database (https://zinc15.docking.org/catalogs/dbfda/; Version: 2018-02-20) (13) as ligands. Prior to docking, the ligand database was converted to a compatible format using Open Babel (v.3.1.1) (14) specifying a split output (‘-m’) and file format (‘-O *.pdbqt’). Resulting docking poses were reviewed with ChimeraX (15). The screening process was repeated with the available Baxα experimental structure (PDB ID: 1F16) to identify non-specific ligands of BaxΔ2.

### 2.2. Cell culture and transfection

Murine hippocampal neuronal cells (HT22; Kerafast, Boston, MA, USA) were maintained in Dulbecco’s Modified Eagle Medium (DMEM) supplemented with 10% fetal bovine serum (FBS). Cells were seeded in 6-well plates and grown to 60-70% confluence. Cells were transfected with BaxΔ2-GFP, Baxα–GFP and GFP using Lipofectamine 3000 (Invitrogen, Waltham, MA, USA), as described previously (10). Transfected cells were cultured for 24-48 hours (as indicated in the text) prior to subsequent analysis.

### 2.3. Drug treatment and cell death assay

HT22 cells were pre-treated with selected drugs (as indicated in the text) for 2 hours prior to transfection. Cells were subsequently incubated for 48 hours before immunostaining or cell death assay. Caspase activity was inhibited by treatment with caspase 8 inhibitor z-IETD-fmk (50 μM) (R&D Systems, Minneapolis, MN, USA) concurrent with transfection. Floating and adherent cells were collected separately following the incubation period. Cell death was calculated by dividing the number of floating GFP-positive cells by the total number of GFP-positive cells.

### 2.4. Immunofluorescent staining and image

Glass cover slips were sterilized and coated with 0.1% gelatin for 30 minutes. Transfected cells were fixed with 4% paraformaldehyde and permeabilized with 0.2% Triton X-100 phosphate-buffered saline (PBS). Coverslips were incubated with primary antibodies, TIA-1 (1:50) and BaxΔ2 (1:200), and Alexa Fluor secondary antibodies (1:200; Invitrogen, Waltham, MA, USA) as described previously (4). Nuclei were stained with DAPI. Fluorescent images were obtained using a Keyence BZ-X710 All-in-One Fluorescence Microscope and analyzed by ImageJ (v1.52d) (16).

### 2.5. Statistics

All experiments were performed at least three times. Statistical significance between groups was determined by student t-test (two-tail) using GraphPad Prism 7.0.0 for Windows (www.graphpad.com; GraphPad Software, Boston, MA, USA). A p-value of < 0.05 was considered statistically significant.

## 3. Results

### 3.1. In silico screening chemical ligands binding to the BaxΔ2 monomer

Because an experimentally determined BaxΔ2 structure is not currently available, we utilized a predicted BaxΔ2 structure for this study. This structure was generated by building a homology model based on Baxα followed by a molecular dynamics (MD) simulation (12). MD simulations show that Baxα retains a conformation consistent with the published crystallographic structure (17). We hypothesize that the exposed hydrophobic pocket of BaxΔ2 – resulting from the missing helix α1 encoded by exon 2 (Fig. 1A) – may allow ligands to bind preventing homo-oligomerization.

**Figure 1.**
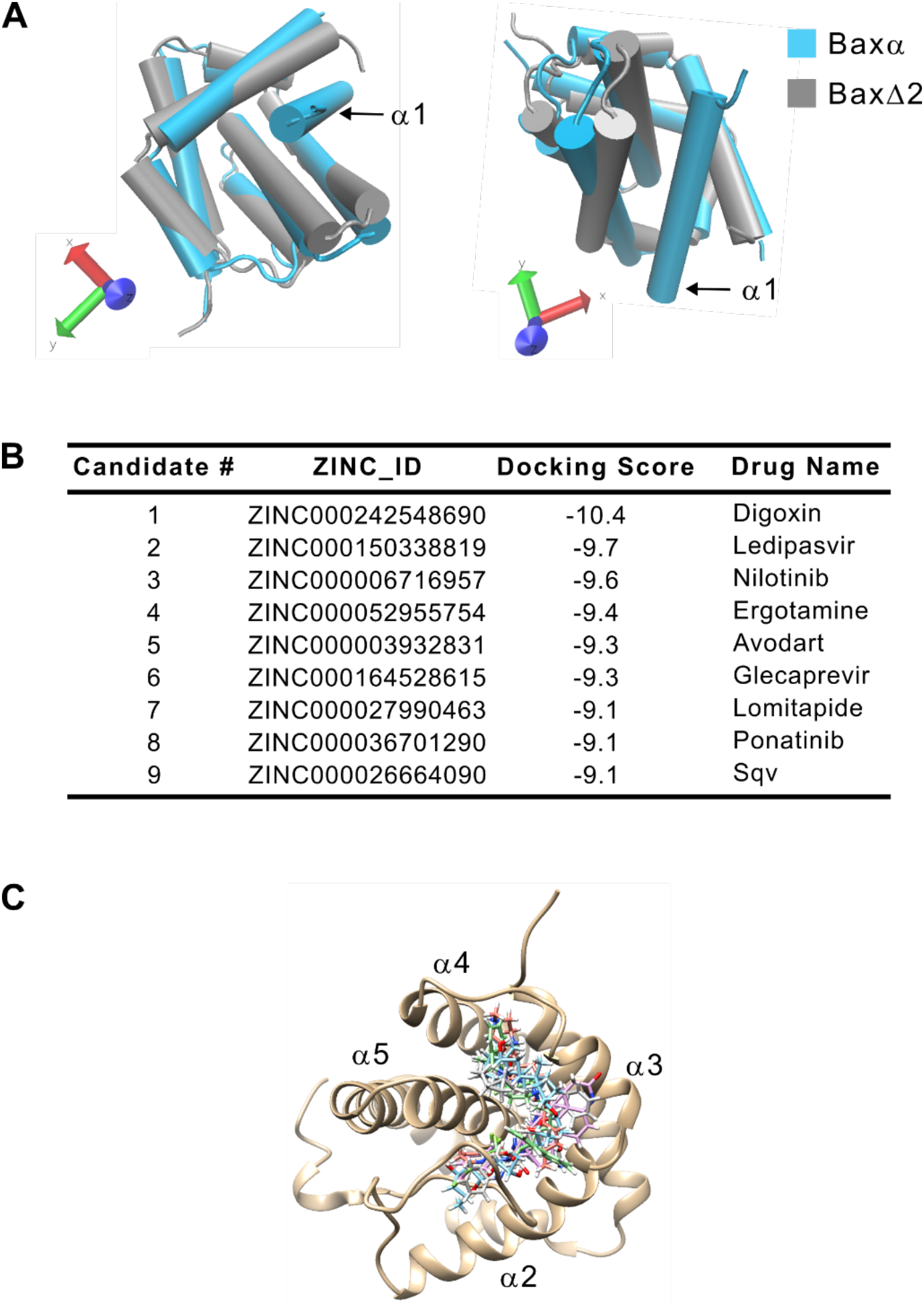
Protein structure prediction and virtual drug screening. (A) Predicted protein structures Baxa and BaxΔ2. (B) Top scored ligands for BaxΔ2 from drug screening using an FDA approved drug bank. (C) Docking poses of top-scored ligands in the modeled BaxΔ2 structure.

Next, we subjected the predicted BaxΔ2 monomer structure to virtual screening of FDA-approved drugs. The nine candidates with the most favorable docking scores for BaxΔ2 are shown in Figure 1B. The same drug library was screened against the Baxα monomer, yeilding a completely different set of ligands (data not shown). All top candidates of BaxΔ2 interacted with the hydrophobic pocket formed by helices α2-5, (Fig. 1C). Given that the hydrophobic pocket is considered responsible for the homo-oligomerization of proteins in the Bcl-2/Bax family (18, 19), and that BaxΔ2 aggregation is required for cell death (10), we hypothesized that binding of candidate ligands to the hydrophobic pocket may prevent BaxΔ2 aggregation, thereby preventing cell death. Furthermore *in silico* analysis showed that the digoxin cardiac glycosides are composed of two structural features, the sugar (glycoside) and the non-sugar (cardiotonic-steroid) moieties (Fig. 2A), with the later proposed to interact with Na-K ATPase (17). The interaction sites with BaxΔ2 are more concentrated in the digoxin glycoside moiety, rather than steroid moiety (Fig. 2B). This could allowing future modification of the drug away from its primary cardio action.

**Figure 2.**
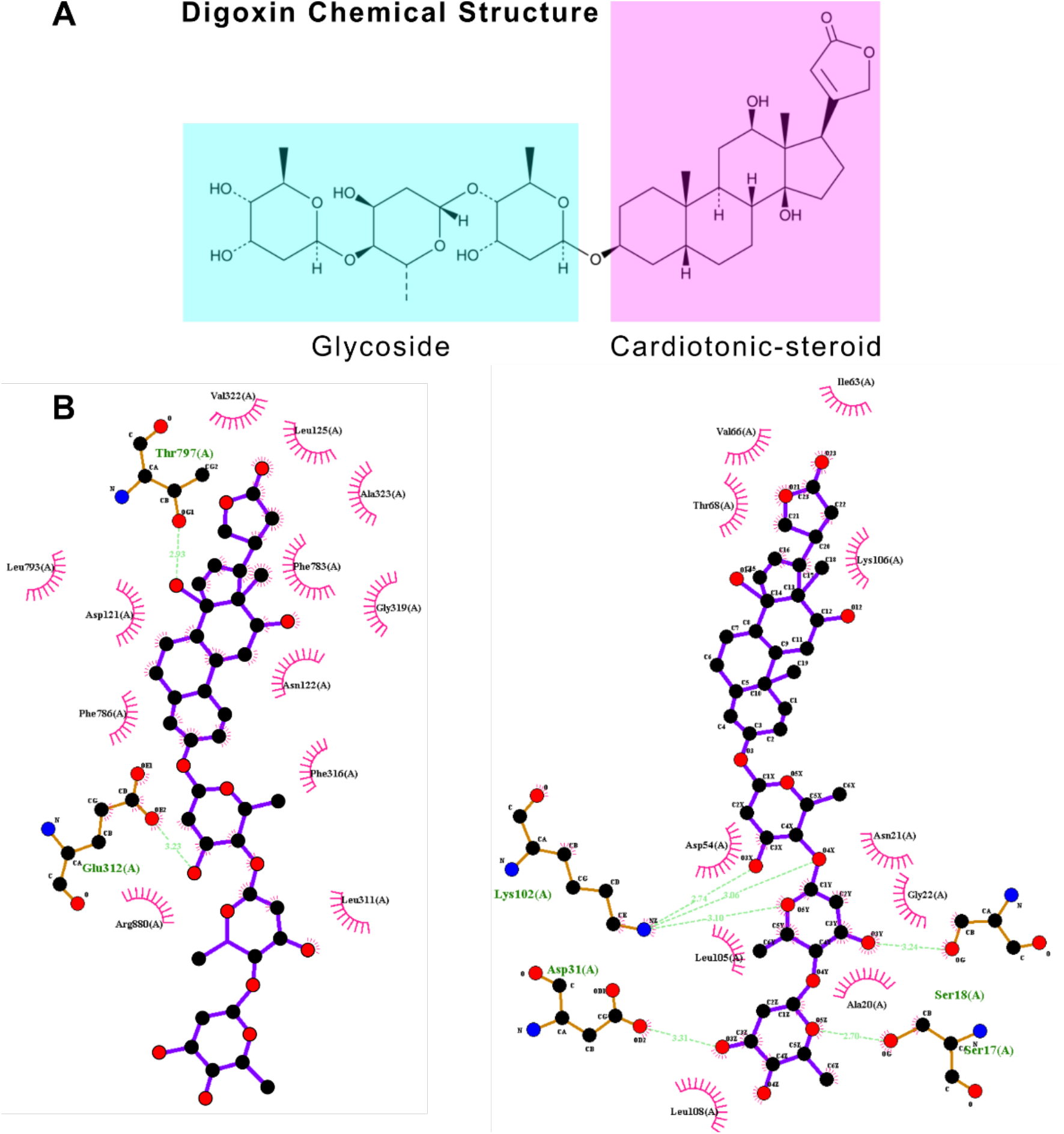
Digoxin structure and protein target residue interactions. (A) Digoxin is a glycoside joined to a cardiotonic-steroid via an ester bond. (B) The interactions of digoxin with its FDA approved target (Na,K-ATPase; Left) are mostly hydrophobic and occur around the cadiotonic-steriod component. The interactions with BaxΔ2 (Right) are a combination of hydrophobic and hydrogen bonds, occurring mostly around the glycoside component. Hydrophobic interactions are depicted by semi-circles with radiating lines. Hydrogen bonds are indicated by dashed lines, and bond lengths are provided.

### 3.2. Digoxin blocks BaxΔ2 aggregates and cell death

To test whether the ligands identified *in silico* could prevent aggregation of BaxΔ2, six candidate ligands were evaluated using an *in cell* system. Mouse neuronal cells (HT22) were used to test ligand cytotoxicity (Fig. 3A). Because HT22 cells do not express endogenous BaxΔ2, we ectopically expressed BaxΔ2-GFP with and without ligand treatment. Cells were monitored periodically under a fluorescence microscope for up to 48 hours. Initial observation indicated that all candidate ligands – except digoxin – exhibited low solubility under our test condition (e.g., ledipasvir) or had no effect on BaxΔ2-induced cell death (e.g., saquinavir and ponatinib) (Fig. 3A). Digoxin showed no cytotoxicity up to 500 nM and reached maximal cell death at 1 uM, therefore, subsequent experiments were conducted using between 50 and 200 nM digoxin.

**Figure 3.**
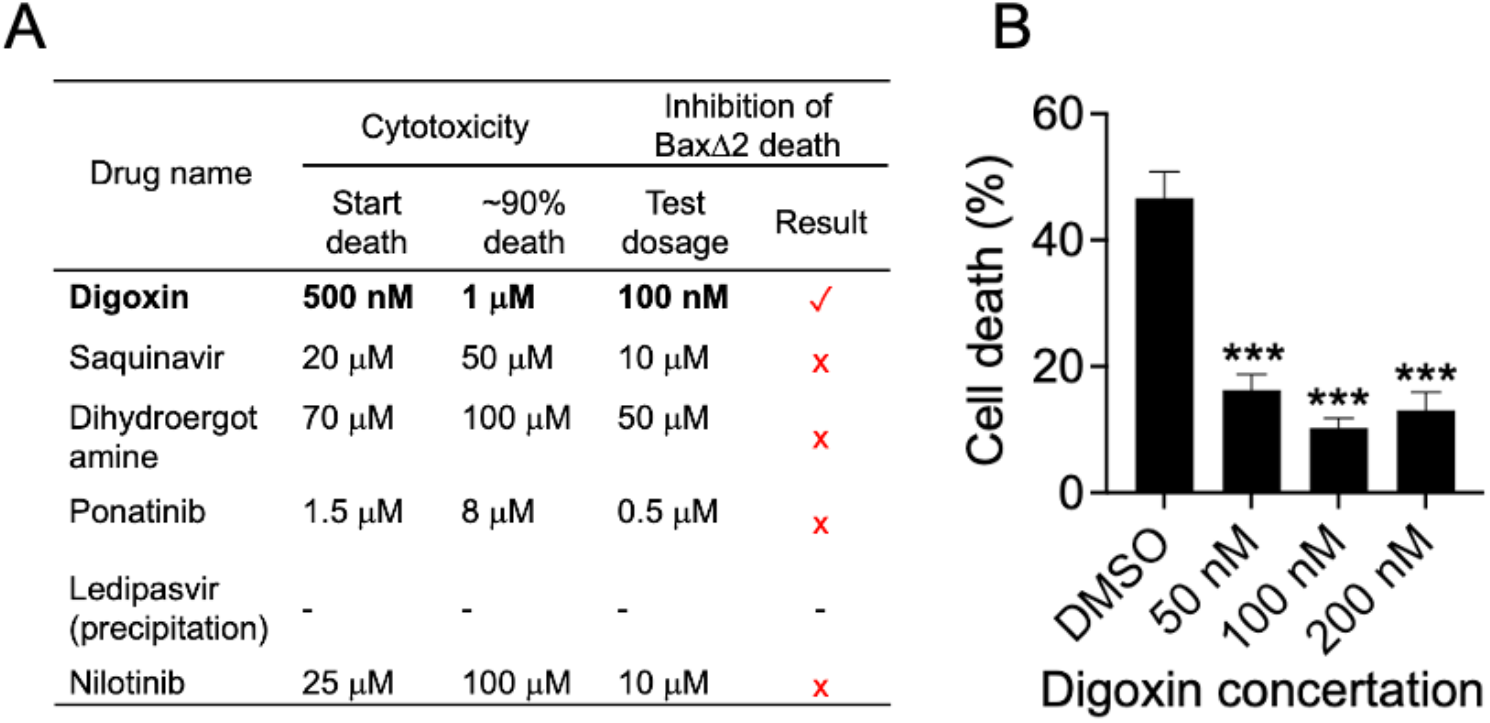
Protective function of digoxin in neuronal cells. (A) Cytotoxicity and cell death inhibition of selective top-scored ligands. For cytotoxicity of drug concentration estimation, HT22 cells were treated with selective drugs at various concentration. The concentrations for the start-point of cell death (5-10%) and at the max cell death (>90%) were recorded. For cell death inhibition estimation, HT22 cells were transfected with BaxΔ2-GFP and the transfected cells were periodically observed under fluorescence microscope up to 48h. “x” indicates no sign of protection or similar to the control without drug treatment. “√” means some visible protection. “-” indicates not tested due to the low solubility under our experimental condition. (B) Cell death assay of digoxin pre-treated HT22 cells transfected for 24h with BaxΔ2-GFP in the presence of different concentrations of digoxin indicated. All values are presented as the mean± standard deviation (SD). The student t test was performed. ***, p< 0.001.

To further evaluate digoxin’s protective function, HT22 cells were pre-treated with varying doses of digoxin (0, 50,100, and 200 nM) and transfected with BaxΔ2-GFP for 24h. Live imaging and quantitative cell death assays showed digoxin could significantly decrease BaxΔ2-induced cell death (Fig. 3B), indicating protection of neuronal cells from BaxΔ2-induced cell death at nanomole concentrations.

### 3.3. Digoxin inhibition is specific to BaxΔ2, and independent of its cardio therapeutic role

To further test the specificity of digoxin for BaxΔ2, we performed hydrogen bond analysis. The result showed that BaxΔ2 helices α2 and α5 could form three hydrogen bonds with digoxin’s sugar chain (Fig. 4A), whereas no hydrogen bonds were found between Baxα and digoxin (Fig. 4B). Additionally, docking scores indicated digoxin interacts more favorably with BaxΔ2 than (-10.9) than it does with Baxα (-8.8). Consistently, the cell death assays showed that digoxin inhibited BaxΔ2-induced, but not Baxα-induced, cell death in HT22 neuronal cells (Fig. 4C). These results suggest that digoxin’s inhibition is specific to BaxΔ2.

**Figure 4.**
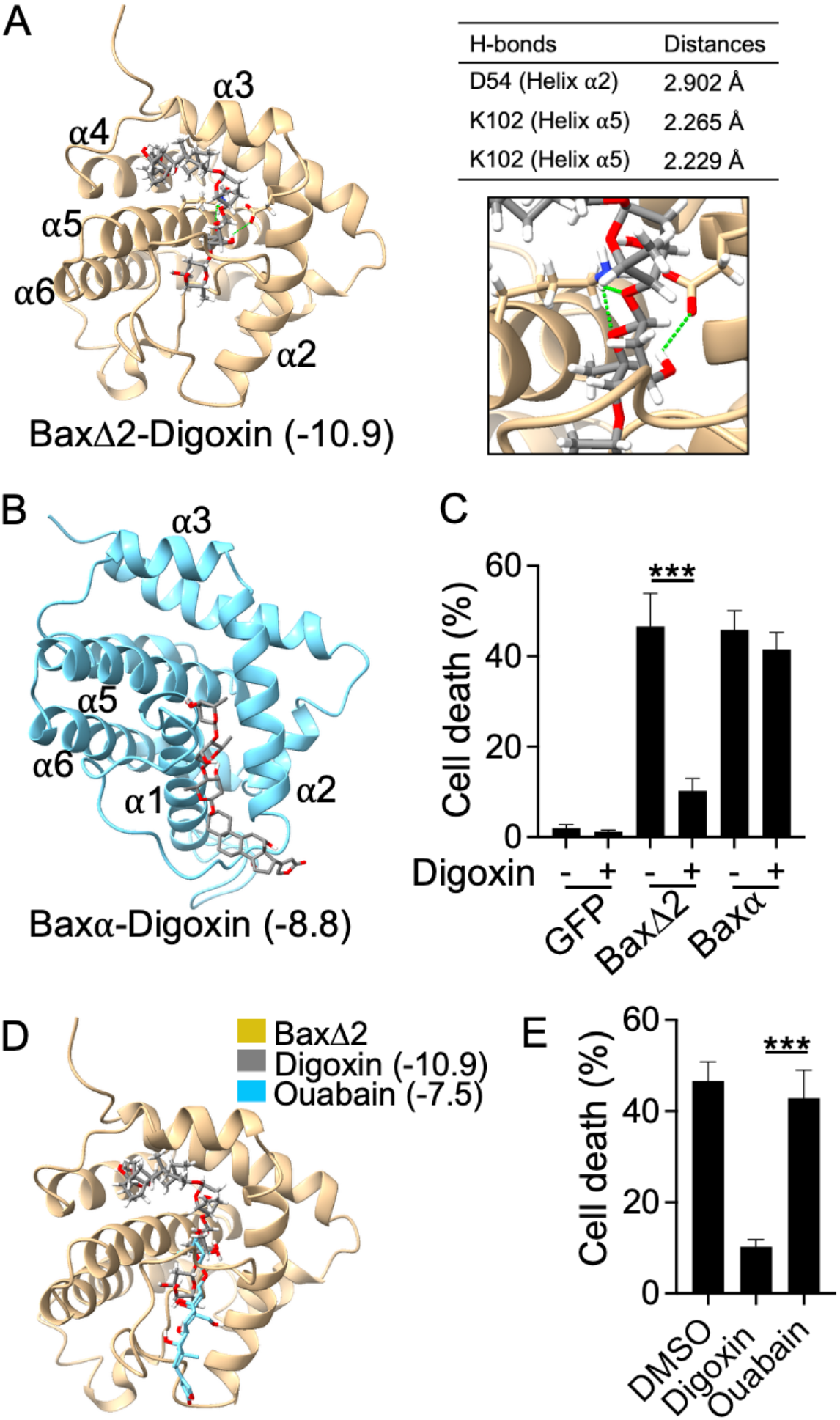
Specificity of digoxin towards to BaxΔ2. (A) Docking pose of BaxΔ2 and digoxin, with hydrogen bonds are marked as green dash line. (B) Docking pose of Baxα and digoxin. (C) Cell death assay of digoxin (200 nM) pre-treated HT22 cells transfected with GFP, BaxΔ2-GFP, or Baxα-GPF for 24h. All values are presented as the mean± standard deviation (SD). (D) Docking pose comparison of digoxin and Ouabain on BaxΔ2, the docking scores are indicated in the parenthesis. (E) Cell death assay of digoxin (200 nM) or Ouabain (200 nM) pre-treated HT22 cells transfected with BaxΔ2-GFP for 24h, DMSO was use the as solve control. The student t test was performed. ***, p< 0.001

Digoxin cardiotherapeutic function is mediated through the inhibition of Na/K^-^ATPase (20, 21). To test whether the effect of digoxin on BaxΔ2 was through the same mechanism, we tested another FDA-approved Na/K-ATPase inhibitor, ouabain, used for treating congestive heart failure (20). Docking results showed that ouabain only interacts with the surface of BaxΔ2, yielding a less favorable docking score compared to that of digoxin (-7.5 vs -10.9, respectively) (Fig. 4D). Indeed, *in cell* death experiments showed that ouabain could not prevent BaxΔ2-induced cell death (Fig. 4E). These results indicate that digoxin’s inhibition of cell death is independent of its FDA-approved cardiotherapeutic function.

### 3.4. Digoxin protection from BaxΔ2-induced cell death is not due to inhibition of BaxΔ2 oligomer formation

To closely monitor the BaxΔ2 protein aggregation process under digoxin treatment, HT22 neuronal cells were transfected with BaxΔ2-GFP for ∼14 hours. By this time point, most transfected cells remained attached to the culture dish, allowing detailed imaging of aggregates. As expected, the even cytosolic distribution of the GFP control was not effected by digoxin treatment (Fig. 5A; top panel). In contrast, BaxΔ2 proteins showed variable cellular distributions: ranging from diffuse without aggregates (Categrory 1), to small and large aggregates (Category 2 and 3, respectively), regardless of digoxin treatment. There is no consistent difference between untreated and treated groups accross all catergories (Fig. 5B). Immunobloting analysis further showed that BaxΔ2 protein levels were lower in the digoxin-treated group (Fig. 5C), but this difference was not statistically significant (Fig. 5D). We were unable to assess BaxΔ2-mediated stress granule formation under these conditions, as transfection with the GFP control alone triggered stress granule formation in our *in cell* model. Together, these data suggest that digoxin-mediated protection from BaxΔ2-induced neuronal cell death is through a mechanism other than the inhibition BaxΔ2 oligomerization.

**Figure 5.**
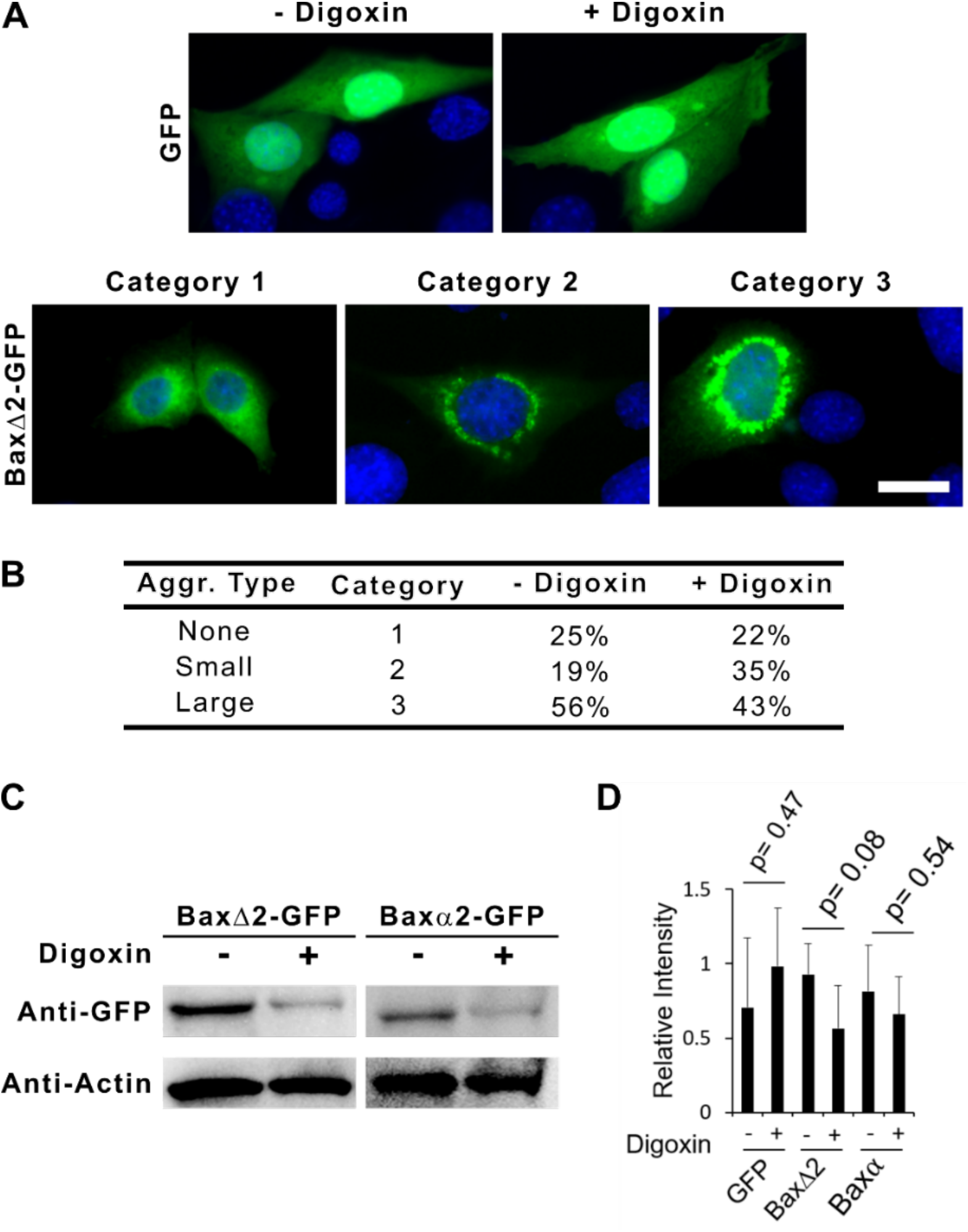
Intracellular localization of BaxΔ2 under digoxin treatment. HT22 cells were pre-treated with digoxin (100 nM) for 2 hours before transfection of GFP or BaxΔ2-GFP for additional 14 hours before imaging. (A) Fluorescent images of cellular distributions of GFP and BaxΔ2-GFP, the latter is classified as Cat1 (no aggregates), Cat 1 (small aggregates) and Cat 3 (large aggregates). (B) quantitation of (A), 50 transfected cells per group were analyzed for each category. (C) Immunoblotting analysis of protein expressions for Baxα-GPF and BaxΔ2-GFP (D) Quantitation of the blots (performed from 3-5 independent experiments).

## 4. Discussion

*In silico* analysis showed that digoxin fits well into the BaxΔ2 hydrophobic pocket, with interaction sites distinct from those present in Na/K-ATPase. Unfortunely, *in cell* results do not support the hypothesis that the digoxin pro-suvival effect is mediated by blocking BaxΔ2 aggregation. Interestingly, digoxin-mediated protection from BaxΔ2-induced neuronal cell death appears to be selective: its protective effect is specific to BaxΔ2-induced cell death and is independent of its primary cardio-action on Na/K-ATPase. It is possible that that digoxin regulates BaxΔ2 protein expression or downstream death pathways. Further investigation is needed to explore the underlying mechanism.

In addition to its well-documented function for cardiac conditions, digoxin has several “off-label” uses related to neuroprotection, neuroinflammation mitigation, and improving cognitive functions in various neurodegenerative disease models (21, 22). For instance, in a dementia rat model, digoxin significantly reduced STZ-induced hippocampal cell death, neuroinflammation, and cholinergic deficiency, ultimately rescuing the rats from memory loss (21). Furthermore, there are endogenous digoxin-like compounds possibly produced by the hypothalamus, adrenal glands, or other tissues (23). This digoxin-like compound is proposed to regulate neuroimmunoendocrine functions and other neurological conditions(24), Although the exact mechanisms of digoxin’s off-label uses and the roles of digoxin-like compounds are unclear, it is evident that digoxin has physiological and therapeutic role far beyond its well-defined cardio-therapeutic effects.

Several limitations exist in this study, and we were unable to confirm *in cell* if digoxin indeed binds to the BaxΔ2 monomer as we predicted. Purificaiton of BaxΔ2 has been unsucessful in using both prokaryotic and eukcaryotic systems due to its toxicity and tendency to aggregate. It is unlikely digoxin can be used directly for treatment of Alzheimer disease due to its narrow therapeutic window and cytotoxicity, however, it may be possible to modify its structure to eleminate this cardiotonic effect while retaining its pro-survival function. Futher investigation may help drug design aimed at preventing protein aggregation in Alzheimer’s disease.

